# ddOTs: a multiplexed quantitative ddPCR approach for resolving overlapping hepatitis B virus transcripts to decipher cccDNA-driven transcription

**DOI:** 10.1101/2025.08.27.672646

**Authors:** Nazim Sarica, Irene Meki, Oceane Lopez, Basile Jay, Gael Petitjean, Christine Neuveut

**Author notes:** present address: Animal Production and Health Laboratory, Joint FAO/IAEA Centre of Nuclear Techniques in Food and Agriculture, International Atomic Energy Agency, Vienna, Austria.

## Abstract

Chronic hepatitis B virus infection persists as a global health crisis, driving life-threatening liver pathologies such as hepatocellular carcinoma. Central to HBV’s resilience is the covalently closed circular DNA (cccDNA), a viral minichromosome that orchestrates viral transcription and sustains infection. Dissecting cccDNA-driven transcription remains a formidable challenge due to the dense overlap of viral open reading frames, which generates RNA transcripts with shared sequences, complicating precise quantification and hindering efforts to unravel HBV’s transcriptional regulation. To address this bottleneck, we developed a multiplexed assay named ddPCR for Overlapping Transcripts (ddOTs) capable of simultaneously quantifying all major HBV RNA species, including splice variants, with unprecedented specificity and sensitivity. This method overcomes current limitations by leveraging the high-resolution power of ddPCR to deconvolute overlapping transcripts, enabling direct interrogation of individual promoter/enhancer activities and RNA stability. By offering a cost-effective, scalable solution for precise RNA profiling in rare biological samples, this breakthrough tool unlocks new avenues for exploring cccDNA biology and accelerating antiviral drug development.

## INTRODUCTION

Despite the existence of a prophylactic vaccine, Hepatitis B virus (HBV) infection remains a global health challenge, with an estimated 260 million individuals chronically infected and at high risk for developing severe liver diseases such as cirrhosis, liver failure or hepatocellular carcinoma. Chronic hepatitis is responsible for over 1 100 000 deaths annually^1^ (WHO, report 2024). While current therapies suppress viral replication and improve liver function, they fail to eradicate the virus and thus cannot eliminate the risk of liver disease development. Therapeutic failure is partly due to the inability of existing drugs to target the viral episomal covalently closed circular DNA (cccDNA) that persists in the nuclei of infected hepatocytes. This approximately 3,2kb cccDNA serves as the template for all viral transcripts including the pregenomic RNA (pgRNA), which is encapsidated in the cytoplasm and retrotranscribed into viral DNA, a key driver of viral persistence. Understanding cccDNA biology, particularly the mechanisms regulating its transcription, is essential for designing novel therapies targeting the HBV reservoir. cccDNA transcriptional activity can be monitored by quantifying HBV RNAs in the cytoplasm of infected cells. Recent findings suggest that HBV RNA species in patient serum may serve as bio-markers for cccDNA persistence and expression during infection and treatment, underscoring the importance of precise HBV RNAs quantification for both basic research and clinical management^2–9^. cccDNA produces 5 major viral RNAs: 1) a 3,5 kb RNA that consists of two separate RNAs of almost similar size the preCore RNA (preC) and pregenomic RNA (pgRNA) refered in the text as preC/pgRNA, 2) a 2,4 kb large surface protein RNA (PreS1), 3) a 2,1 kb middle and small surface protein RNAs (PreS2 and S), 4) a 0,7 kb X protein RNA (HBx), and 5) splice variants, with Sp1 being the most expressed^10–14^. Due to its small size, the entire viral genome is coding. HBV transcripts are overlapping either completely or partially and share a common polyadenylation site and 3’end (Fig. 1), complicating individual RNA quantification. Current methods face limitations^2,15,16^. Reverse transcription quantitative PCR (RT-qPCR) can distinguish the 3.5-kb preC/pgRNA but requires sequential amplification and subtraction steps prone to primer bias^15^. Northern blotting allows semi-quantification of the main HBV transcripts but lacks sensitivity and requires large quantities of RNA. 5’RACE excludes certain transcripts and is not fully quantitative^2^. Finally, long read sequencing coupled with a capture step is promising but may miss short RNAs, requires substantial input material, and is costly. Thus, none of these technics allow a rapid, clear and sensitive quantification of the different HBV transcripts in a single reliable experiment.

**Figure 1:**
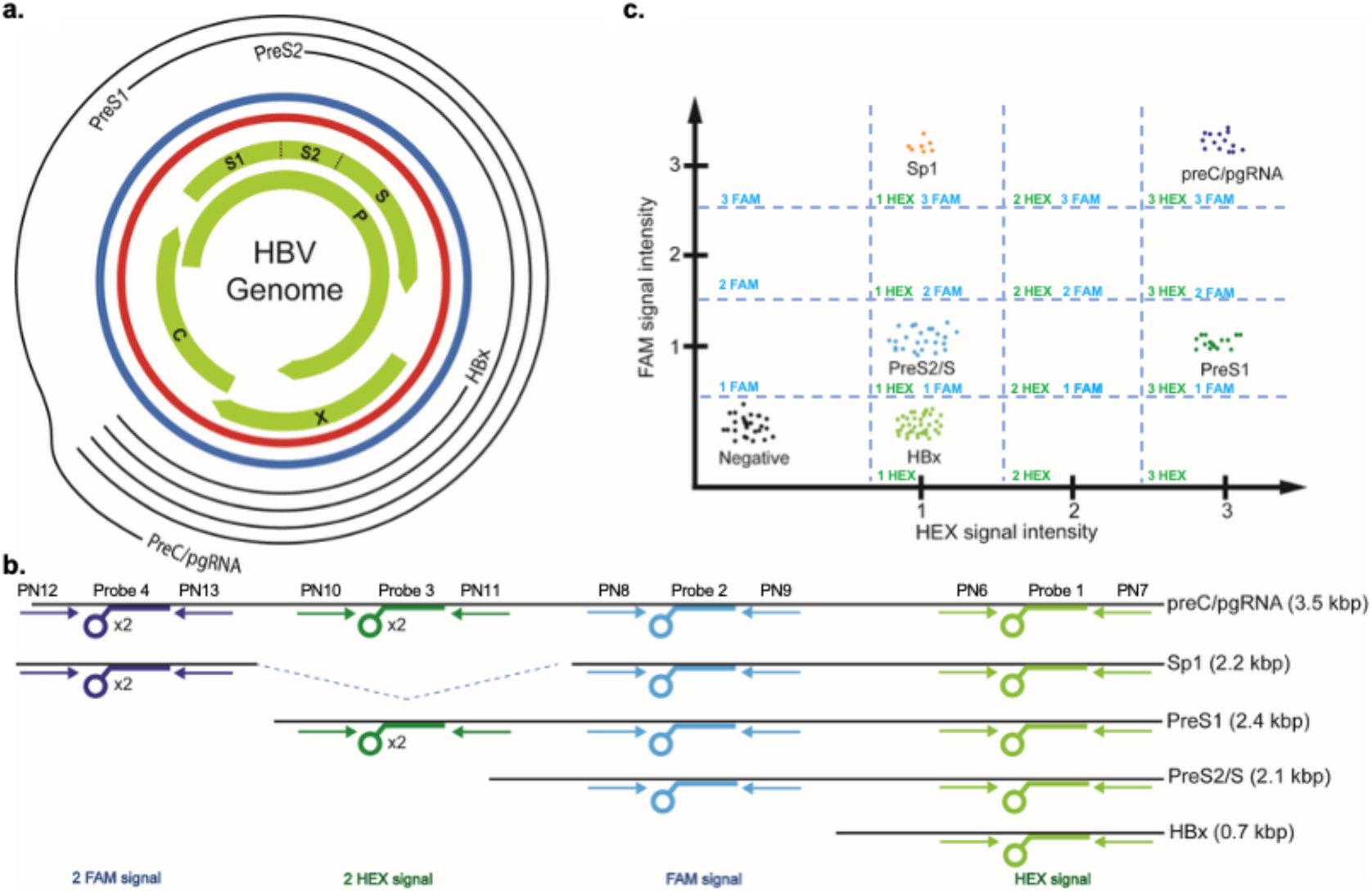
Schematic representation of ddOTs. (a) Schematic representation of the HBV genome with the main ORFs and the HBV RNA transcripts (b) Schematic representation of primers and probe locations on the 5 targeted HBV mRNA species. (c)Theoretical visualization on 2D-scatterplot of ddPCR results using the primers and probes represented in B. X-axis represent HEX signals, Y-axis represent FAM signals. Black dots represent signal-empty droplets. Each coloured dot corresponds to a droplet giving a positive signal for the indicated RNA. Probe 4 and probe 3 concentrations are doubled compared to the two other probes, giving more FAM or HEX signal intensity and inducing a shift on the 2D ddPCR plot.

To address these challenges, we developed a multiplex droplet digital PCR (ddPCR) assay named ddPCR for Overlapping Transcripts (ddOTs) for simultaneous and precise quantification of all major HBV RNA species (HBx, PreS2/S, PreS1, preC/pgRNA, and Sp1) during infection. ddPCR provides absolute quantification with higher precision and lower technical variability than qPCR ^17,18^, and has proven effective in pathogen diagnostics. We validated our method using synthetic HBV-like DNA templates to ensure specificity and accuracy, then applied it to quantify viral RNA dynamics in HBV-expressing cellular models including primary human hepatocytes.

## MATERIALS & METHODS

### Cell culture, HBV production and infection

HepG2-NTCP cells were grown in DMEM-Glutamax supplemented with 10% FBS and 1% Penicillin/Streptomycin^39^. Primary human hepatocytes were maintained in PHH medium (Corning, reference 355056, hepatocyte culture media kit, 500 ml) according to the manufacturer’s recommendations. HepAD38 cells are derived from HepG2 cells and contain an integrated HBV genome (subtype ayw) under tetracycline control^19^. HepAD38 cells were grown in DMEM F12 + Glutamax supplemented with 10% FBS, 1% Penicillin Streptomycin, Hydroxycortisone (0,2µg/mL) and Insulin (5µg/mL). When notified, 0,3µg/mL of Doxycycline was added. HepG2 H1.3Δx cells are also derived from HepG2 cells and contain an integrated 1.3 HBV genome that carries a stop codon mutation in both HBx open reading frames^27^. HepG2 H1.3Δx cells were maintained in Williams E medium.

WT and X-HBV genotype D particles were concentrated from the clarified supernatant of HepAD38 and HepG2 H1.3Δx cells respectively. Supernatants were collected after 5 days in Williams E production medium supplemented with 2% DMSO and digested with Roche DNase I for 1h at 37°C. Nucleocapsids were then concentrated by ultracentrifugation at 32 000xg for 4h on 20% sucrose cushion^27^. Titers of enveloped DNA-containing viral particles were determined by immunoprecipitation with an anti-preS1 antibody (gift from C. Sureau) followed by DNA extraction and qPCR. For infection, only enveloped DNA-containing viral particles (vp) were considered to determine the multiplicity of infection (MOI). PHH and HepG2-NTCP cells were infected by spinoculation (1000xg for 30min) at a multiplicity of infection of 100 vge/cell in respective culture media supplemented with 4% PEG (polyethylene glycol 8000) and 3% DMSO. Medium containing virus was removed the next day, cells were washed with PBS and cultured in medium supplemented with 2% DMSO until processing.

For kinetic analysis, cells were spinoculated for 30min, incubated at 37°C for1h30min, washed with PBS multiple times, and corresponding media complemented with 2% DMSO was added until RNA extraction. For time-point 0, cells were covered with medium containing virus at 100 vge/cell for 1min and then washed multiple times with PBS. Myrcludex was used at 500nM.

For RNA interference, HepG2-NTCP cells were transfected with gene-specific siRNA at a concentration of 10nM using Lipofectamine™ RNAiMAX Transfection Reagent according to manufacturer’s recommendations. PreS targeting siRNA sequence is 5’-CUUCCUAUUAACAGGCCUATT, and control non-targeting siRNA sequence is 5’-UGGUUUACAUGUCGACUAATT.

### Primers and probe design

Primers and probes design were based on HBV genotype D ayw subtype.

Primers used for PCR amplifications and ddPCR were designed using the online Primer3plus software (https://primer3plus.com/Primers). Primers were designed with special attention to the following details: (i) amplicon size between 60 and 200 bp, (ii) primers GC content between 50 and 60%, (iii) avoid repeats of Gs or Cs longer than 3 bases, (iv) when possible 3’ ends with a G or a C.

Probes for ddPCR were designed using IDT PrimerQuest online web interface (https://www.idtdna.com/Primerquest/). The probes design adhered to the following details: (i) probe must not overlap with the prime sequences, (ii) Tm should be 3-10°C higher than that of the primers, (iii) the probe should anneal to the strand that contains more Gs than Cs, (iv) the length of the probe should be less than 30 nucleotides, (v) absence of a G at the 5’ end. Probes were labeled with either 5’ FAM or 5’HEX fluorophores.

Primers PN6, PN7 and Probe 1 (referred as primers/probe set 1, Supplemental Table 1) were used for the amplification and detection of a region spanning HBx ORF. Primers PN8, PN9 and Probe 2 (primers/probe set 2) were used for the amplification and detection of a region spanning PreS2/S ORF. Primers PN10, PN11 and Probe 3 (primers/probe set 3) were used for the amplification and quantification a region spanning PreS1 ORF. Primers PN12, PN13 and Probe 4 (primers/probe set 4) were used for the amplification and quantification a region spanning preC/pgRNA ORF (Fig.1).

### HBV DNA fragments

DNA fragments corresponding to the 4 HBV ORFs preC/pgRNA, PreS1, PreS2/S and HBx were synthetized by PCR using the pFC80 plasmid as template. This plasmid carries HBV genomes cloned head-to-tail into the EcoRI site of pBR322^40^. HBx DNA fragment was generated using primers PN1 and PN5. PreS2DNA fragment was generated using PN2 and PN5. PreS1DNA fragment was generated using PN3 and PN5. pgDNA fragment covering preC/pgRNA ORF was generated using primers PN4 and PN5. PCR products were purified by gel extraction using Monarch Gel Extraction Kit following manufacturer’s protocol. Vortex steps were replaced by gentle pipetting to avoid DNA shearing. DNA were quantified by Nanodrop (ThermoFisher Nanodrop One).

### RNA extraction

Total RNA was prepared using RNeasy Plus Mini Kit (QIAGEN). Manufacturer’s protocol was modified to conserve RNA integrity at maximum. All vortexing steps were replaced by gentle pipetting, and all centrifugations were done at 4°C and 7,000xg with soft acceleration/breaking. Samples were kept on ice throughout the process.

Briefly, 8 x 105 cells were resuspended in 400µL of cold RLT Plus Buffer (supplemented with 40mM DTT) and then mixed with 150µL of QIAGEN RNA Protect Tissue Reagent (QIAGEN ID.76104). The mixture was then transferred to the gDNA eliminator column and centrifuged. 400µL of cold 70% EtOH were added to the flow-through and mixed gently by pipetting. Approximatively 750µL of the mix were transferred to RNeasy Spin Column and centrifuged. Subsequent steps were done following supplier’s protocol. To remove any DNA contamination, samples were treated with 2 units of Invitrogen Turbo DNase for 30 minutes at 37°C followed by addition of DNase inactivation reagent.

### RNA Quality Control

To ensure the accuracy of downstream ddPCR quantifications, quantity, quality, and integrity analysis of extracted RNAs were performed. Samples were processed using Agilent RNA 6000 Nano Kit to measure the RNA Integrity Number (RIN) using Agilent 2100 BioAnalyzer (Agilent Technologies) according to manufacturer’s protocol. Only samples presenting a RIN superior to 10 were conserved for downstream applications.

### Reverse transcription

Total RNA (500ng) was retrotranscribed using ThermoFischer SuperScript IV enzyme and HBV primer PN14 that binds in the 3’ region of HBx. For Rhot2 RNA quantification, total RNA was retrotranscribed using oligo(d)T. Resulting cDNA was then diluted and processed for ddPCR.

### ddOTs

In order to detect the different HBV RNA species using multiplexed ddPCR, we designed four pairs of primers and 4 probes that cover the four ORFs in the viral genome and allow for the discrimination and quantification within droplets via multiplexed PCR of the different cDNA species generated from the different HBV RNA species (Figure 1a, b). Briefly, HBx cDNA hybridizes with primers/probe set 1, giving only one HEX signal in quadrant 1 HEX/0 FAM, while cDNA from PreS2/S transcript hybridizes with 2 primers/probe sets (set 1 and 2) labelled respectively by HEX and FAM appearing in quadrant 1 HEX/1 FAM, cDNA from PreS1 is amplified simultaneously by 3 primers/probe sets (1,2 and 3) in droplets appearing then in quadrant 3 HEX/1 FAM and preC/pgRNA cDNA is amplified simultaneously by 4 primer/probe sets in droplets appearing in quadrant 3 HEX/3 FAM. cDNA synthetized from Sp1 HBV RNA is amplified by primers/probe sets 1,2 and 4 (quadrant 1 HEX/3 FAM).

Droplets with no signal are represented as “negative” on the 2D projection (Fig.1, quadrant 0). Every other signal is considered as positive. ddPCR is a method for absolute quantification of nucleic acid concentrations through the combination of limiting dilution, end-point PCR and Poisson statistics. It is therefore important to do measurements in the right range of positive to negative ratio. Indeed, for high concentrations of positive droplets (>20% of total droplets), probability of having more than 1 molecule per droplet increases exponentially. This phenomenon can bias the measurement by overestimating the number of longer molecules, especially pgRNA molecules in our case (as example, a droplet containing Sp1 and PreS1 molecules will be detected as pgRNA). For low concentrations of positive droplets, most of the partitions will contain zero copies of the target molecules and nearly all positive partitions will contain only one copy of the target molecule. To ensure that almost all droplets contain only one molecule and the measurement remains accurate in multiplex conditions, we recommend keeping the positive to negative ratio lesser than 0,2. Samples must be diluted prior to droplet generation in order to obtain less than 20% of positive droplets.

As RNA integrity is primordial for discrimination and quantification of the different HBV RNA species by RT-ddPCR, we recommend assessing RNA Integrity using Agilent 2100 BioAnalyzer and use RNA presenting RIN over 9.8. In this study, all the experiments were performed using RNA samples with a RIN of 10.

To show the effects of low RNA quality on quantification, we voluntarily induced degradation of RNA sample by heating/vortexing and quantified RNAs by ddPCR (Supplemental Figure 3). On the left, the sample has a RIN of 10, on the right, the sample has a RIN of 5. Measurements show a 3-fold decrease of pgRNA and a 4-fold increase of HBx RNA, confirming the importance of sample quality.

In an amplitude-based multiplex assay, targets can be detected with single dye probes used at different final concentrations. This strategy allows for measurement of multiple targets per reaction and greater separation between signals coming from probes with same dye. In our case, 4 targets are being quantified in a tetraplex reaction. HEX-probes 1 and 3 are used at 250 and 500nM respectively, and FAM-probes 2 and 4 are used at 250 and 500nM respectively^41^. With these experimental conditions, probe 1 hybridization to its target will give the equivalent of 1 HEX signal while probe 3 hybridization will generate the equivalent of 2 HEX signals^41^. Similarly, probe 2 hybridization will give 1 FAM signal while probe 4 hybridization will give 2 FAM signals. These settings offer a clearer separation of clouds and facilitate the detection of unwanted/misplaced signals (Fig. 1c; 2a) due to degradation/shearing of RNAs. As example, a 3’ degraded PreS1 RNA losing the binding regions for Probe 1 and 2 will appear in the “2 HEX” quadrant instead of contaminating HBx measurements in “1 HEX” quadrant. Finally, these different relative concentrations will also allow better detection of splice variants in HBV expressing cells.

A variety of experimental settings were used for the development of ddOTs. RNAs are extracted from 4 x 10^5^ cells using RNeasy Plus Mini Kit from QIAGEN with precautions to avoid RNA shearing. RT-ddPCR is then performed using a two-step approach. cDNAs are first synthetized using Superscript IV enzyme and then used as template for droplet generation and measurements using Bio-Rad Multiplex ddPCR kit. We do not recommend the use of the Bio-Rad One-Step ddPCR kit as our results (data not shown) unveiled that the RT has limited ability to amplify simultaneously multiple fragments inside the same droplet.

ddPCR was performed using ddPCR Multiplex Supermix according to the manufacturer’s protocol (Bio-Rad). Controls containing DNase-treated samples without RT were systematically ran in parallel to confirm effectiveness of DNase degradation. Briefly, 5µl of diluted cDNA were mixed with 5µL 4× ddPCR Multiplex Supermix, 1,8 µM of primers, 250 or 500 nM of each probe, 0.27µL of DTT 300mM, in a total reaction volume of 20 μL. The mixture was then partitioned into 20,000 droplets using the Automated Droplet Generator. PCR amplification was performed in a C1000 Touch thermal cycler (Bio-Rad, Hercules, CA, USA) with the following amplification program: 5 min at 95 °C, 40 cycles of denaturation for 30 s at 94 °C and annealing for 60 s at 59 °C (ramping rate set to 2 °C/s), final incubation step for 10 min at 98 °C, and kept at 4°C until fluorescence measurement. Fluorescence was measured on QX200 or QX600 Droplet Digital PCR Systems. Measurements were treated using QX Manager software.

### ddPCR data analysis

After fluorescence measurements in a QX200 (or QX600) Droplet Reader (Bio-Rad), the data were analyzed using QX Manager Software 2.3 Standard Edition (Bio-Rad), which automatically calculated absolute sample concentration. Fluorescence amplitude threshold to distinguish positive from negative droplets was based on amplification of negative controls (water, no reverse transcriptase, and non-HBV infected samples). Althought not required, quantification of Rhot2 expression may be used for normalization in cases where the number of replicates is low or sample quality is variable.

## RESULTS

### Validation of ddOTs using DNA fragments covering the main HBV ORFs

To specifically quantify each of the overlapped HBV RNAs, we developed a multiplexed ddPCR approach. We designed four pairs of primers/probes sets targeting the 4 main HBV ORFs (HBx, PreS2/S, PreS1 and preC/pgRNA) (Fig. 1b). Following reverse-transcription of RNA and partitioning of cDNA into droplets, individual cDNA molecules are simultaneous analyzed with the 4 primer/probe sets, generating a specific fluorescence profile (Fig.1c). To validate the HBV multiplexed ddPCR strategy, we first used individual purified DNA fragments spanning the 4 HBV ORFs. By testing known concentrations of each DNA fragment individually, we confirmed accurate amplification, quantification, and fluorescence profiles (Supplemental Fig. 1). Evidenced by droplets outside expected clusters, shearing occurred at low levels (Supplemental Fig. 1). Terminal dilution assays demonstrated linear quantification (R^2^=0,9989) and sensitivity down to 0.4 copies/µL (Supplemental Fig. 2). We validated the assay using a mixture containing equal amounts (10 molecules/µL) of all 4 HBV DNA fragments. The 2D projection (Fig. 2) shows distinct clusters for each HBV DNA, with negative droplets labeled accordingly. Absolute quantification confirmed the specificity and accuracy of the primer/probe sets, with <6% of positive droplets corresponding to sheared DNA (Fig. 2; Supplemental Fig. 1).

**Figure 2:**
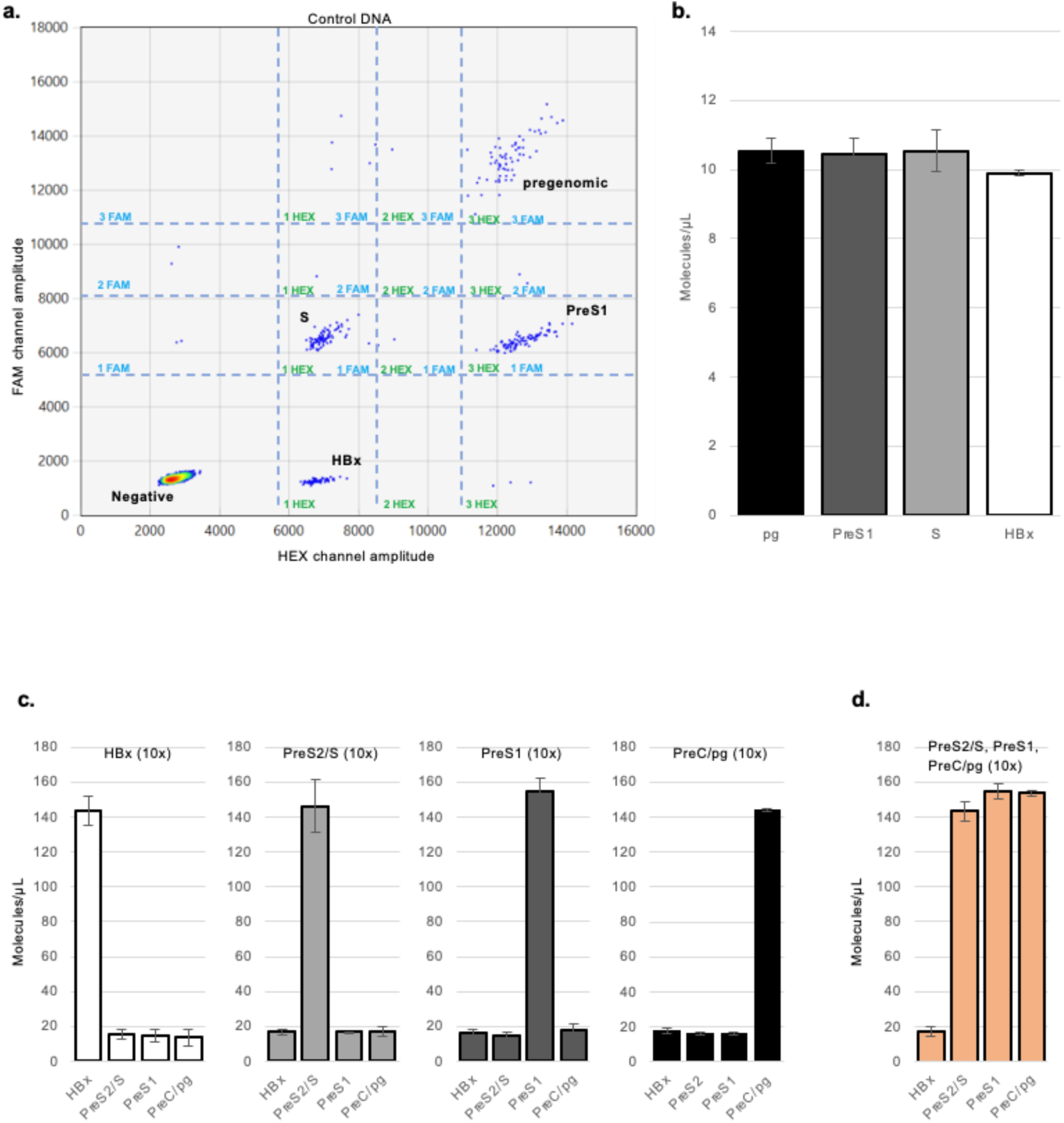
Validation of ddOTs quantification and specificity using synthetic DNAs fragments covering the 4 HBV ORFs. (a) to (b) Indicated PCR-amplified and purified HBV fragments were mixed at a concentration of 10 molecules/µl each and then quantified using ddOTs. (a) 2D ddPCR plot of synthetic DNA fragments HBx, PreS2, PreS1, and pregenomic quantification on QX Manager Software. X-axis represents HEX signal amplitude; Y-axis represents FAM signal amplitude. Clouds corresponding to the different HBV fragments are indicated. Sparse unannotated dots correspond to sheared molecules. Negative droplets are indicated (b) Absolute quantification of each HBV fragments present in samples measured by ddOTs. Errors bars represent SD of three independent experiments. (c) Quantification by multiplexed ddPCR of the 4 HBV DNA fragments present at different concentration. As indicated on each graph, DNA fragments were mixed at a concentration of 15 molecules/µL each excepted for the indicated fragment added at a concentration of 150 molecules/µl (10 times more concentrated). Errors bars represent SD of three independent experiments. (d) Same as in (c) except that all the DNA fragments are mixed at a concentration of 150 molecules/µl each except HBx DNA fragment added at a concentration of 15 molecules/µL. Errors bars represent SD of three independent experiments.

To assess performance under imbalanced conditions, we tested samples where one DNA fragment was 10-fold more concentrated than the others. The multiplexed ddPCR accurately quantified all targets regardless of overrepresentation (Fig.2c). Similarly, when HBx DNA was underrepresented, all species were reliably measured (Fig.2d). Altogether, the multiplexed ddPCR assay robustly quantifies HBV DNA fragments corresponding to major ORFs.

### Analysis of HBV RNA expression in HepAD38 cells by ddOTs

Having established the conditions for the multiplexed ddPCR, we validated the assay using HBV RNA expressing cells. HepAD38 cells are derived from the HepG2 hepatoma cell line and contain an integrated cDNA copy of the HBV pgRNA under tetracycline-controlled expression. The promoter region comprises a minimal CMV promoter (−53 relative to the start site) and heptamerized upstream tet-operators. In the absence of tetracycline, this promoter drives a strong pgRNA expression, while its activity, and to a lesser extend the expression of other HBV transcripts, is silenced upon tetracycline treatment^19^. We first quantified HBV RNA molecules in untreated HepAD38 cells using multiplexed ddPCR (Fig. 3a, 3c). The 2D representation in figure 3a shows the expression levels of the 4 main HBV RNAs: HBx, PreS2/S, PreS1, and preC/pgRNA, with the latter being the most abundant, consistent with other studies^19,20^. Notably, the Sp1 splice variant was detected at relatively high levels, representing 14% of total HBV transcripts here. This aligns with reports that Sp1 constitutes up to 17% of HBV transcripts ^10–14,21,22^. Additional clusters observed in the assay may correspond to other spliced HBV RNA species, which are known to vary in abundance across cell lines ^23^. While RNA degradation or shearing could partially explain these extra clusters, the combined signal from the four main RNAs and Sp1 accounted for ∼95% of the total, leaving only 5% attributable to minor splice variants or fragmented RNA.

**Figure 3:**
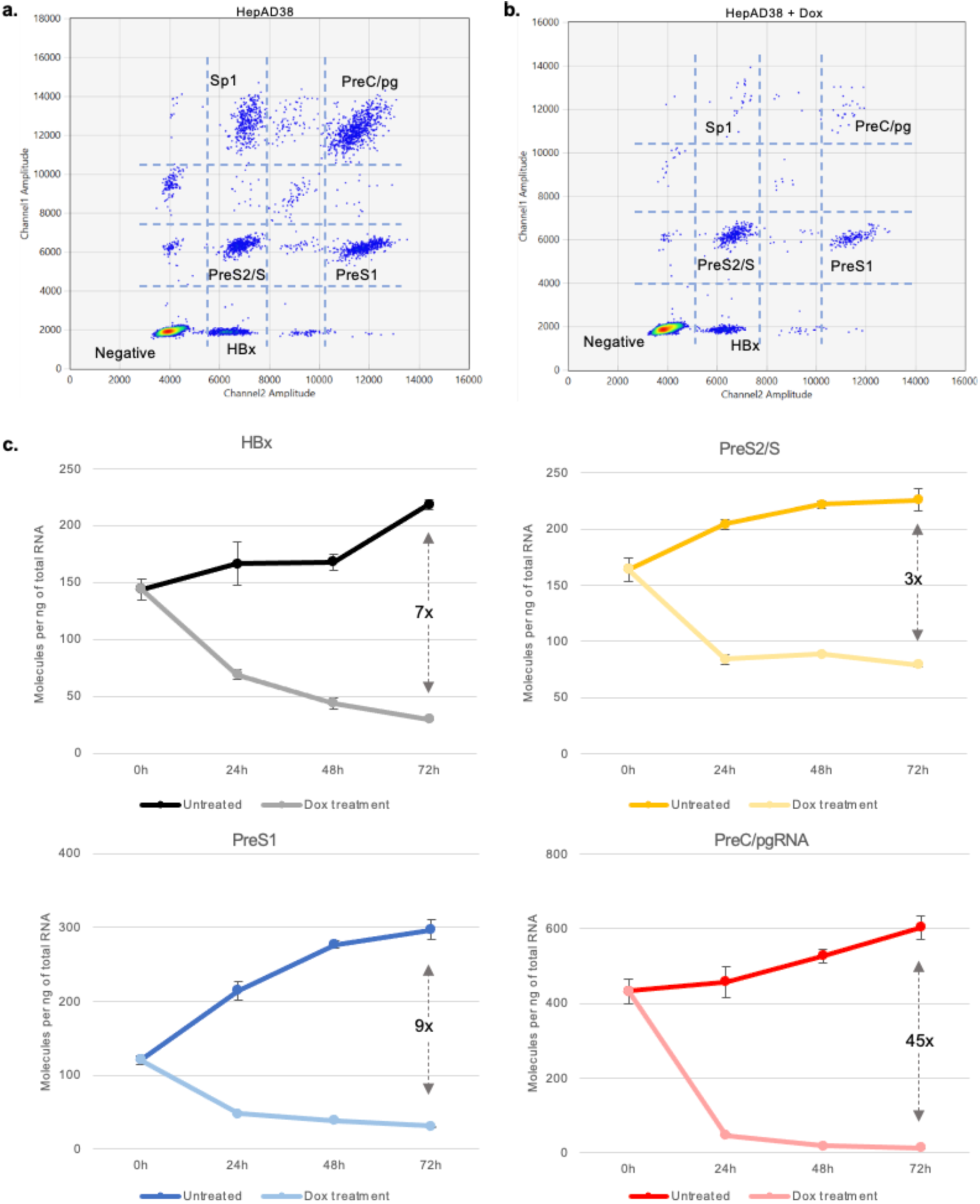
Quantification of HBV RNAs levels in HepAD38 cells treated or not with Doxycycline. HBV RNAs expression in HepAD38 cells has been quantified by ddOTs over 72-hours time-course experiment following treatment with or without Doxycycline (0,3µg/mL). 0h corresponds to the time when the cells were plated with or without Doxycycline treatment. Total RNA was extracted at different time after plating and quantified by ddOTs. (a) Visualization of the 2-D ddPCR plot showing the detection of each HBV RNAs in non-treated HepAD38 cells 48h after plating. (b) 2-D ddPCR plot showing the detection of each HBV RNAs in HepAD38 cells treated with Doxycycline for 48h. (c) Quantification of the indicated HBV RNAs level at the indicated time after plating using ddOTs in HepaAD38 cells treated or not with doxycycline. Results are expressed as the number of copies of indicated RNA/ng of total RNA. Mean ± SD of 3 experiments is shown.

To evaluate the assay-specificity for detecting oriented transcriptional changes, we analyzed HBV RNAs expression in HepAD38 cells after silencing the tet-CMV promoter with doxycycline. HBV RNA levels were quantified at multiple time points post-treatment. Fig. 3b and 4c reveal a sharp decline in pgRNA and Sp1 levels within 24 hours of doxycycline exposure. HBx, PreS2/S, and PreS1 RNAs also decreased, albeit to a much lesser extent. Although these genes are regulated by their own promoters, the tet operator’s repressive effect may propagate across the HBV genome, as previously suggested^20^. However, residual expression of HBx, PreS2/S, and PreS1 RNAs might originate from cccDNA transcription, which accumulates in HepAD38 cells cultured without tetracycline. In conclusion, our multiplexed ddPCR assay robustly detects specific variations in HBV RNA expression, enabling precise tracking of transcriptional dynamics.

**Figure 4:**
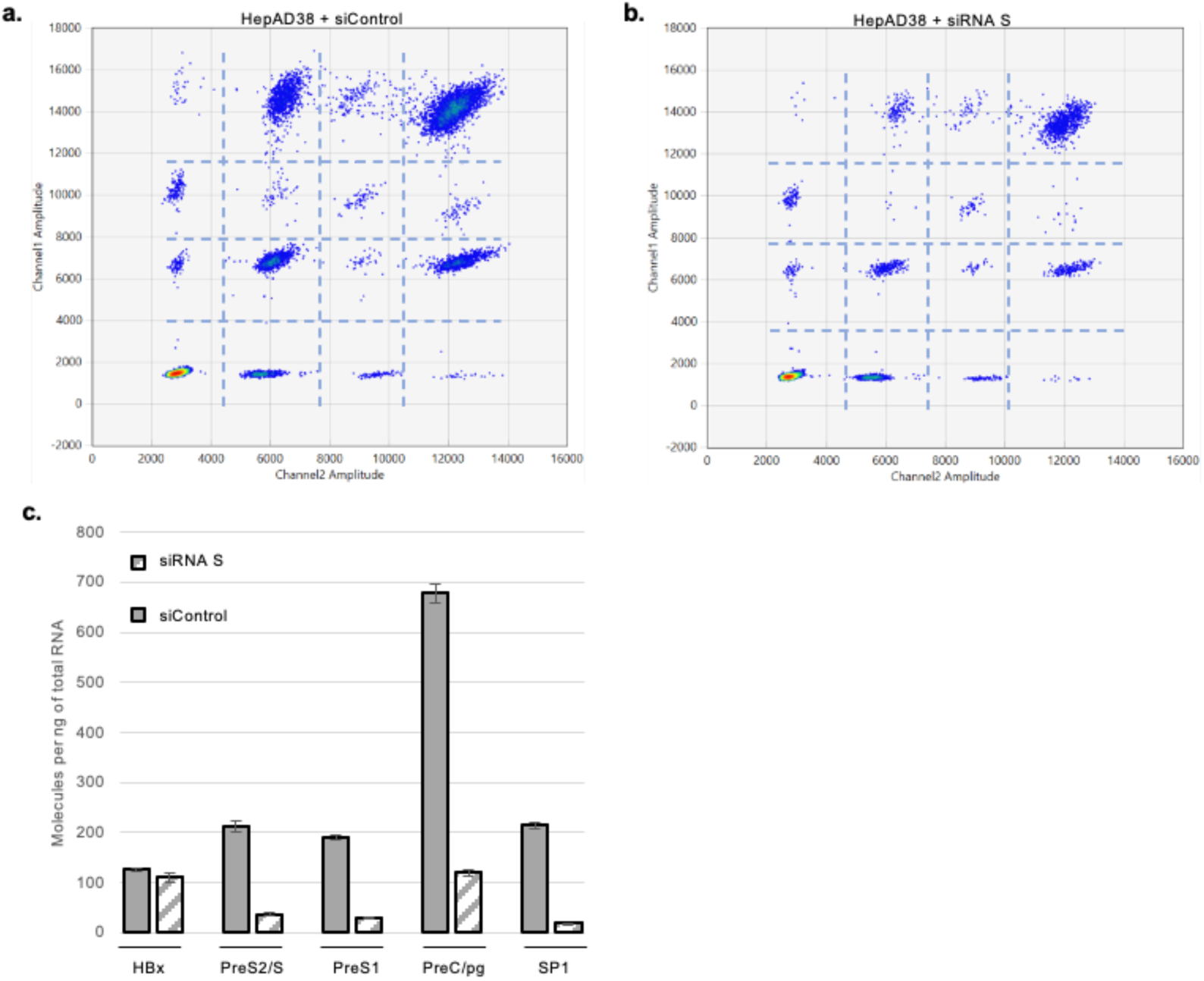
HBV RNA levels quantification in HepAD38 cells transfected with siRNA targeting the S region. HepAD38 cells were transfected with an siRNA targeting a region in S ORF that overlaps with preC/pgRNA, Sp1, PreS1 and Pres2/S HBV RNAs or with a control siRNA. 72 hours after transfection, HBV RNA levels were quantified by ddOTs. (a) (b) 2-D plot of ddPCR showing the detection of the different HBV RNAs in HepAD38 cells transfected with control siRNA (a) or with the siRNA targeting the S region (b). (c) Quantification of the number of copies of the indicated HBV RNAs in HepAD38 cells transfected with the siRNA S or with siControl by ddOTs. Errors bars represent the SD of three independent experiments.

### Analysis of HBx RNA expression level in HepaD38 cells by ddOTs

Depending on the study and the technique used to assess HBV mRNAs expression, the level of HBx mRNA in HBV replicating cells varies widely, ranging from negligible to as high as PreS2 mRNA levels^2,4,15,24–26^. Our results suggested that in HepAD38 cells, HBx mRNA levels are comparable to PreS2/S or preS1 mRNA levels. Since HBV RNA quantification relies on reverse transcription of viral RNAs followed by amplification via ddPCR, one may question whether incomplete reverse transcription of longer, more abundant HBV mRNAs (e.g., PreS2 or preC/pgRNA) could generate shorter cDNAs that will be recognized and quantified as HBx mRNA. Similarly, degradation of longer HBV mRNAs may lead to erroneous quantification of HBx mRNAs. To rule this out, we measured the level of HBx RNA levels in HepAD38 cells with or without siRNA-mediated silencing of all HBV transcript except HBx. The siRNA targeted the S ORF region. As shown in Figure 4, siRNA targeting the S region induced a ∼80% reduction in PreS2/S, PreS1, preC/pgRNA, and Sp1 mRNA levels, while HBx RNA levels remained unchanged. These results validate the ddOTs approach for absolute quantification of all HBV RNAs, including HBx RNA.

### Analysis of HBV RNA expression during infection using ddOTs

The HBx protein has been shown to be required for HBV cccDNA transcription^27–29^. HBx acts primarly by degrading the SMC5/6 complex, which silences HBV transcription via a mechanism that is not yet fully understood but results in the establishment of a repressed chromatin state. A recent publication suggests that in the absence of HBx, cccDNA can still transcribe HBx mRNA^26^. To assess whether HBV promoters were differentially affected in absence of HBx, we analyzed the expression of HBV mRNAs in HepG2-NTCP cells and primary human hepatocytes (PHH) infected with wild type (HBV WT) or HBV deficient for HBx expression (HBV X-).

As shown in Figure 5 a-c, in the presence of HBx, all HBV mRNAs were detected, with preC/pgRNA and PreS2/S being the most highly expressed. As observed in HepAD38 cells, the Sp1 splice variant was also detected in relatively high quantities. In HepG2-NTCP cells infected with HBV X-, the levels of all HBV RNA species decrease. Specifically, preC/pgRNA and PreS1 levels decrease by 3.9-fold, whereas preS2/S and HBx RNAs decrease by 2.3-fold and 1.6-fold, respectively. In infected PHH (Fig. 5d-f), all HBV mRNAs and Sp1 splice variant were again detected, with PreS2/S remaining the most highly expressed. In PHH infected with HBV X-, transcript levels decreases compared to those infected with HBV wt: preC/pgRNA and Sp1 decrease by 7-fold, PreS1 by 6-fold, PreS2/S by 5-fold, and HBx by 2-fold. In both cell lines, Sp1 expression decreased proportionally to the reduction in total preC/pgRNA plus Sp1 suggesting that transcriptional repression imposed by SMC5/6 does not impact splicing.

**Figure 5:**
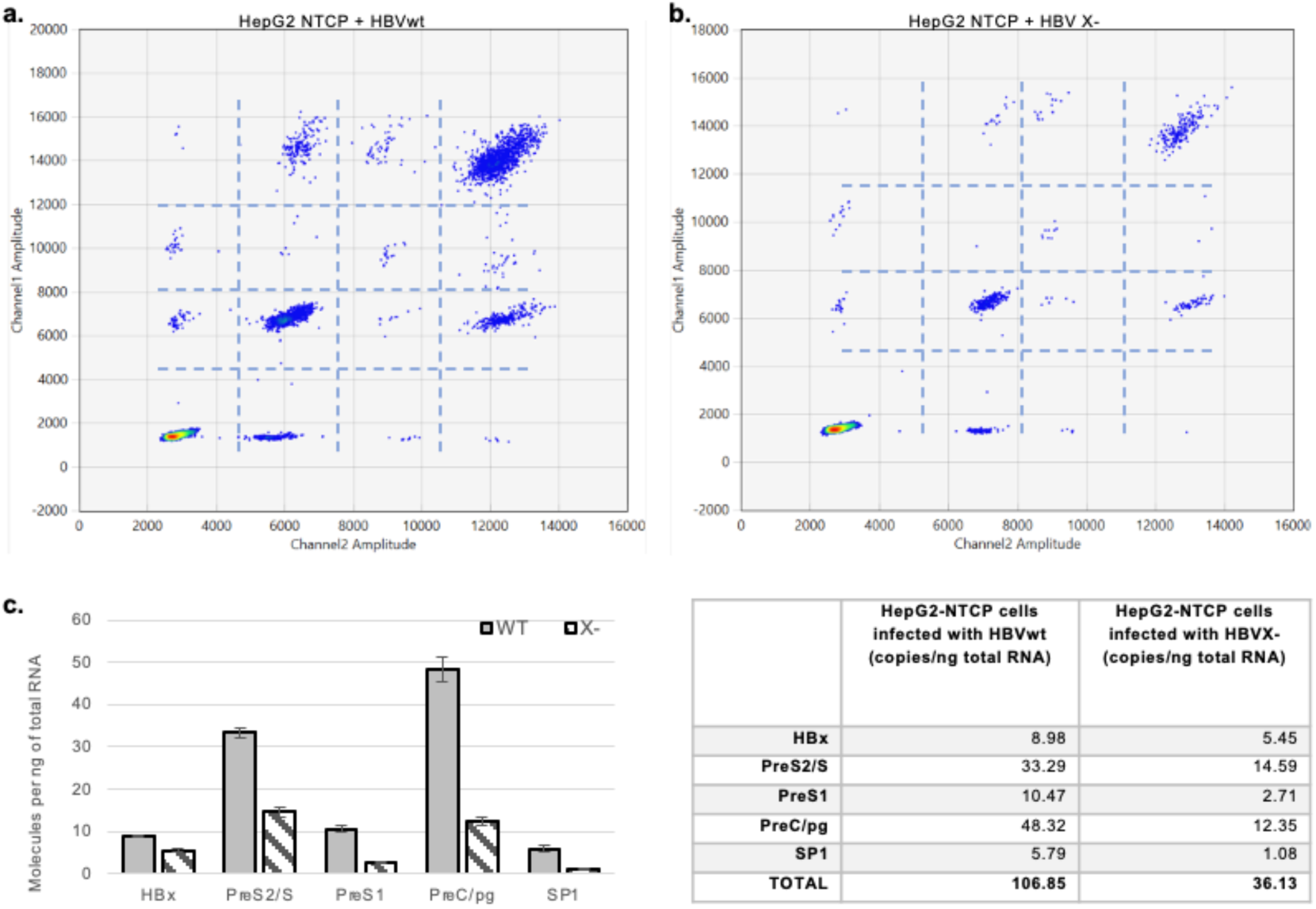
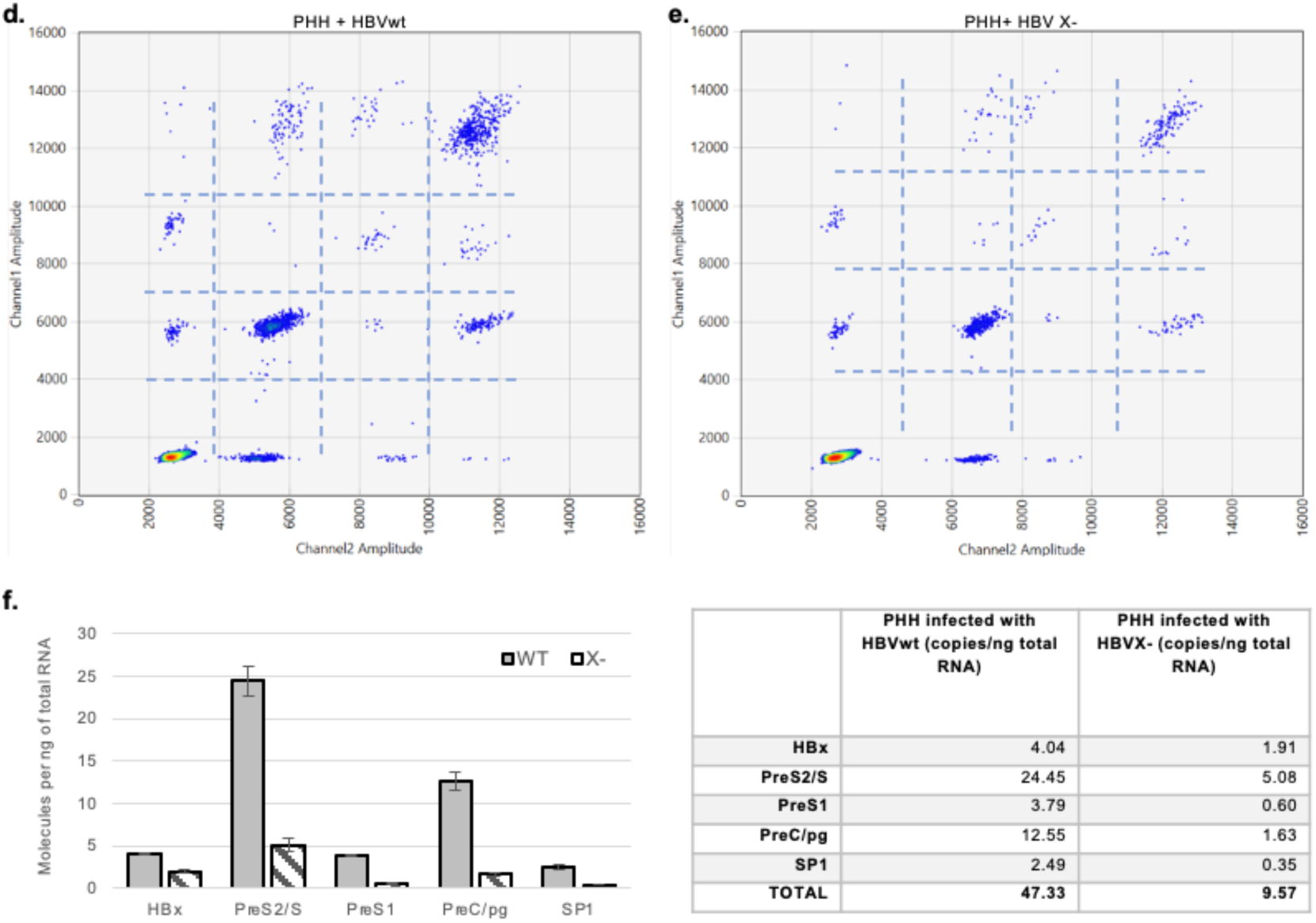
HBV RNA levels in HepG2-NTCP cells or human primary hepatocytes infected with WT or X-HBV. (a) to (c) HepG2-NTCP cells were infected with WT or X – HBV at a MOI of 100 vp/cells. Total RNA was collected 6 dpi and HBV RNAs were quantified by ddOTs. (a) (b) 2D ddPCR plot of HBV RNAs analysis in HepG2-NTCP cells infected with HBV wt (a) or HBV X-(b). X-axis represents Channel 2 amplitude corresponding to HEX signals, Y-axis represents Channel 1 amplitude corresponding to FAM signals. (c) Absolute quantification of the different HBV RNA species in cells infected by WT or X-viruses. Results were normalized to the number of cells and expressed as number of copies per ng of total RNA. Errors bars represent the SD of three independent experiments. (d) to (f). PHH were infected with WT or X – HBV at a MOI of 100 vp/cells. Total RNA was collected 5 dpi and HBV RNAs were quantified by ddOTs. (d) (e) 2D ddPCR plot of HBV RNAs analysis in PHH cells infected with HBV wt (d) or HBV X-(d). X-axis represents Channel 2 amplitude corresponding to HEX signals, Y-axis represents Channel 1 amplitude corresponding to FAM signals. (f) Absolute quantification of the different HBV RNA species in cells infected by WT or X-viruses. Results were normalized to the number of cells and expressed as number of copies per ng of total RNA. Errors bars represent the SD of three independent experiments.

Overall, our results validate the use of ddOTs to quantify HBV transcription in the context of infection and suggests that HBV promoters are differentially affected in the absence of the HBx protein.

### Analysis of dynamics of HBV RNA expression in early stages of infection by ddOTs

The dynamics of HBV RNA expression during early steps of infection remains unclear. Studies suggest that HBx RNA is present in the incoming viral particle and is translated, leading to the degradation of smc5/6 and consequently initiating transcription from cccDNA, while other studies propose that HBx RNA is expressed before other HBV RNAs in order to enable cccDNA transcription^30,31^. Finally, Lucifora and colleagues proposed that repression is established shortly after cccDNA transcription begins, allowing the expression of HBV RNAs including HBx RNA^27^. To investigate the kinetics of HBV RNA expression and determine whether HBx RNA is an early transcript, we infected HepG2-NTCP cells with HBV WT or HBV X-viruses and quantified HBV RNA levels at different time-points post-infection (Fig. 6a). HBx RNA is detected as early as 2 hours post-infection (h.p.i) with a slight increase from 4h to 12 h.p.i, coinciding with the initial detection of preC/pgRNAs. A pronounced increase in HBx RNA occurred at 16 h.p.i., simultaneous with the detection of all HBV RNA species (Fig. 6a, 12–16 h.p.i.). Between 16–24 h.p.i., all HBV RNAs showed a steady increase, though preC/pgRNA, preS2, and preS1 RNAs exhibited far greater amplification (33-fold in average), compared to HBx RNA increase (5-fold) from 24 to 48 h.p.i. In HBV X-infected cells, HBx RNAs were detected early but remained constant until 16 h.p.i.

**Figure 6:**
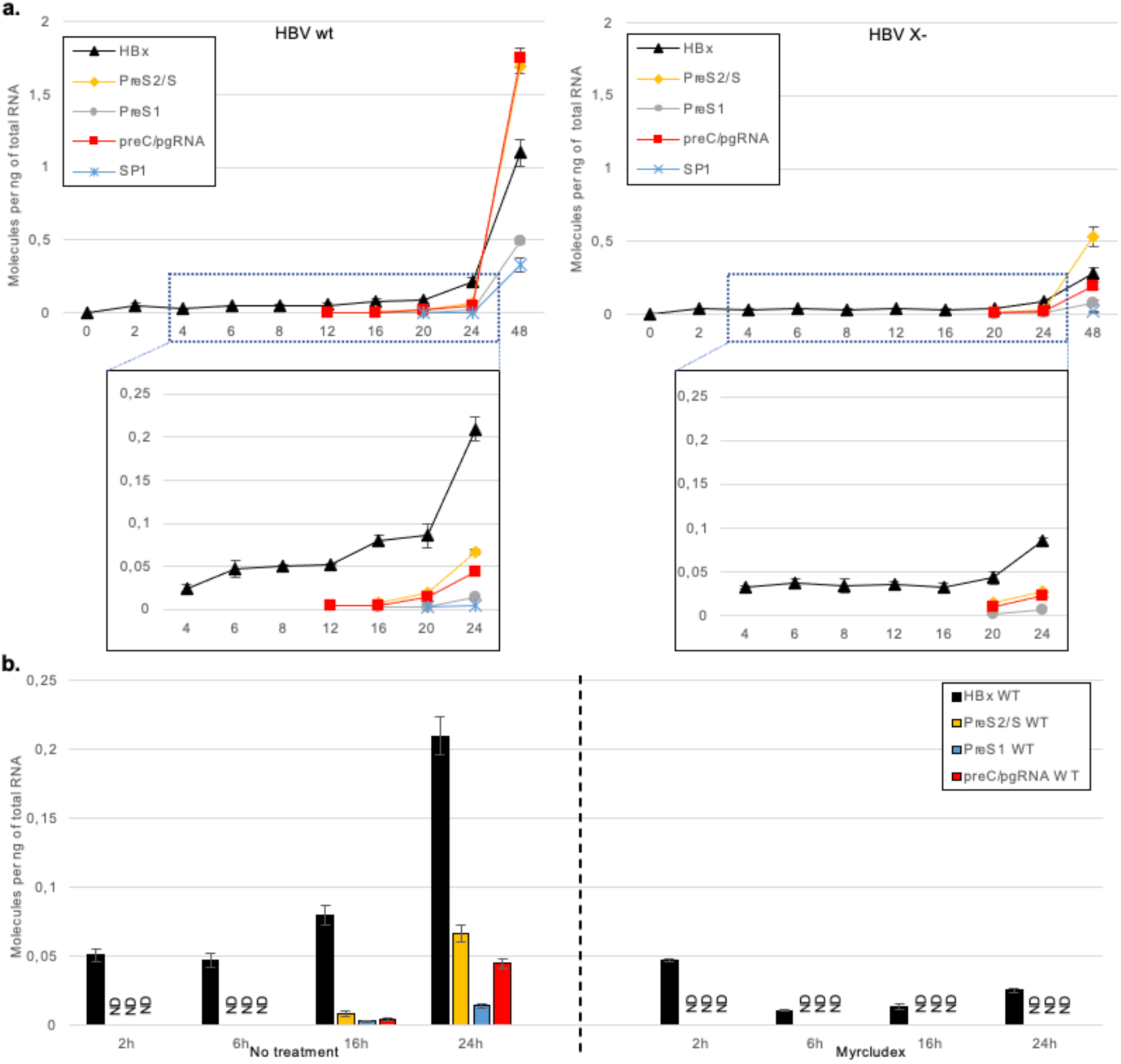
Analysis of HBV RNAs kinetic in HepG2-NTCP cells using ddOTs. HepG2-NTCP cells were infected with WT or X – HBV at a MOI of 100 MOI vp/cells and total RNA was extracted at the indicated time after infection. (a) Absolute quantification of the 4 main HBV RNAs and Sp1 in cells infected with WT (left) or X-(right) viruses. (b) Absolute quantification of the 4 main HBV RNAs and Sp1 in HepG2-NTCP cells treated or not for 2 hours with 500nM Myrcludex and infected with HBV wt at a MOI of 100 MOI vp/cells. Results are expressed as number of copies per ng of total RNA. Errors bars represent the SD of three independent experiments.

All HBV RNA species emerged concomitantly at 20 h.p.i., coinciding with a rise in HBx RNA. Unlike HBV WT, HBV X− transcription remained low over time, confirming HBx’s necessity for sustained cccDNA transcription. Notably, while HBV RNA levels were similar between WT and X− viruses at transcriptional onset, HBV X− transcription was delayed (20 h.p.i. vs. 16 h.p.i.) and failed to amplify over time. Furthermore, HBV X− infection skewed RNA abundance: preS2/S RNA dominated, unlike WT infections where preC/pgRNA and preS2/S RNA were predominant, suggesting that HBV promoters are regulated differentially in response of the cellular environment.

To test if HBx transcripts detected during the first hours after infection originate from inoculum-derived enveloped particles, we pretreated HepG2-NTCP cells with myrcludex, which block the NTCP receptor, 2 hours before infection. In treated cells, HBx RNAs decreased rapidly by 2 h.p.i., with no subsequent viral transcripts detected. In contrast, untreated cells showed rising HBx RNA levels by 2 h.p.i., indicative of neo-transcribed RNA (Fig. 6b). This suggests a portion of early HBx RNA enters cells via NTCP-independent routes ^2,32^, while subsequent transcription requires NTCP-mediated entry.

Altogether, these results validate ddOTs for sensitive quantification of HBV transcripts, even at low-abundance stages. They confirm HBx RNA’s early detection, its essential role in sustaining cccDNA transcription, and differential regulation of HBV promoters. While incoming HBx RNA is detectable, its functional significance in HBV replication warrants further study.

## DISCUSSION

The compact structure of the HBV genome and the overlap of different open reading frames have been a long-time roadblock for the development of a quantitative affordable daily usage assay for understanding HBV transcription dynamics. In this study, we developed the ddOTs method to quantify the major HBV RNAs: preC/pgRNA, PreS1, PreS2/S, HBx and Sp1. Using DNA covering the four HBV ORFs we first showed that we were able to quantify accurately each of the DNA species even under imbalanced conditions. We next validated this approach using different model of HBV RNA expression. Using ddOTs we were able to achieve absolute quantification of the 4 main HBV RNAs (preC/pg, PreS1, PreS2/S, HBx) as well as the splice variant Sp1. Using siRNAs targeting the S region and thus decreasing the expression of preC/pgRNA, PreS1, PreS2/S and Sp1 RNAs, we showed that ddOTs allows the specific quantification of the HBV X RNAs. Finally using different settings known to modulate differentially HBV transcription (ie treatment of HepaD38 cells with doxycycline or infection of HepG2-NTCP cells or PHH with HBV X-virus) or performing time course analysis, we demonstrated that ddOTs can be used to accurately quantify oriented transcriptional changes.

Using terminal dilution assays, we showed that ddOTs can detect as few as 0,4 copie/µL using DNA fragments covering the HBV ORFs. It will thus be possible to use ddOTs to analyze HBV transcription in limited materials such as patient’s biopsies. However, our technic’s efficiency is directly dependent of the quality of the RNA and may require standardized tissue-banking methods allowing high quality RNA extraction. Moreover, our method has been developed for the detection of genotype D HBV RNAs and should thus be further adapted for the detection of the different HBV genotypes. The usability of degenerate primers for multiplex ddPCR amplifications has not been investigated yet and may be an easy solution for detection of HBV RNA of any genotype.

HBx RNA expression level reported in the literature is highly variable depending on the technics used to detect/quantify HBV RNAs. To further demonstrate that ddOTs specifically quantify HBx RNA and not degradation products of larger HBV RNAs, we induced the degradation of all the HBV RNA except HBx using an siRNA targeting the S ORF and we showed that the level of HBx remain constant. Our results are in line with previous studies showing that HBx mRNA is expressed at a level comparable to those of structural genes and remains relatively constant over time^2,26,31,33^. Discrepancy with other studies may come from technical caveat as the removal of short transcripts during sample preparation for long read sequencing or the variation in PCR amplification of the different HBV regions when using a subtractive based RT-PCR assay^15,16^

Using ddOTs we were able to clearly identify and quantify simultaneously the 4 main HBV transcripts and Sp1 splice variant in HepAD38, HepG2-NTCP cells and PHH. The relative abundance of each RNA species is in line with previous observations, i.e. preC/pg RNA being reported as the more abundant HBV transcripts in HepG2-NTCP cells and PreS2/S in PHH^26^. Sp1 is the most abundant splice variant in every cell line we tested, which is also consistent with published results^10–13^. Beside Sp1, we were also able to identify additional clouds that could represent the different spliced RNAs described in the literature^23,34,35^. Recent studies suggest that spliced HBV RNA can represent up to 30% of total RNAs. Lim *et al.*^23^ showed a high spectrum of splice variants present in HBV-infected cells and patients by analyzing more than 500 RNA-seq libraries. However, we cannot completely rule out that these additional clouds may in part be due to HBV RNA degradation or shearing. The development of probe with new fluorochromes will help to the identification of spliced variant. To this day, limitations of Bio-Rad QX Manager software do not permit to quantify molecules simultaneously bound by more than 3 different fluorochromes. Once this software limitation is addressed, design of new primers/probe sets with different fluorescence will allow better separation and discrimination between transcripts, such as PreS2 and S or between splice variants. ddOTs methodology may thus be applied to other compact-genome systems that present overlapping transcript sequences, such as prokaryotes, viruses, and eukaryotes^36–38^.

The relevance of ddOTs to study the transcriptional regulation of the HBV promoters in the context of the full HBV genome is first supported by our results in HepaD38 cells that contain a cDNA copy of the HBV pgRNA. Using ddOTs we were able to quantify simultaneously and independently the expression of the 4 main HBV RNAs and of Sp1 in cells treated or not with doxycycline over a period of 72h. We clearly observed a differential sensitivity to doxycycline treatment of the different promoters with a strong decrease of more than 45-fold for the HBV pgRNA directly controlled by the tet CMV promoter compared to preS1, preS2/S and HBx RNAs that are regulated by their own promoter. In a more physiologically relevant system, we were able to assess the regulation of the 4 main HBV promoters individually using HepG2-NTCP cells or PHH infected by HBV wt or HBV X-viruses. In accordance with the literature, the expression of HBV RNAs was strongly repressed in the absence of HBx expression. However, our results suggest that the HBV promoters do not respond similarly to smc5/6 silencing. Recently Peng and collaborators, using 5’-terminal single-cell sequencing analysis, showed that while HBV RNAs expression is strongly decreased in human primary hepatocytes infected by HBV X-, some HBV RNAs are still expressed, albeit at low level, and the majority of them correspond to long HBx RNA starting at a non-conventional TSS in the enhancer 1^26^. These results suggest that HBV promoters are differentially regulated. In the line with these findings, recent results from Prescott and collaborators show that HBx RNA expression strictly relies on chromatin assembly at HBx promoter compared to the other HBV promoters. While regulatory elements present in the HBV promoters and enhancers have been well characterized, little is known yet on their regulation in the context of the cccDNA in the cell. ddOTs is thus a promising tool to study HBV transcriptional and post-transcriptional regulation.

Using ddOTs we assessed the kinetics of HBV RNAs expression and showed that HBx RNAs are detectable very early after infection. While part of this HBx RNAs, as suggested by others, are delivered from the inoculum, they however seem to enter the cell through an NTCP-independent mechanism. Moreover, their level decreased quite rapidly over time (6h post infection), as shown in myrcludex treated cells. One may ask the significance and the role of these incoming HBx RNA in virus replication. It has been suggested that they serve as the template for the translation of HBx proteins allowing cccDNA transcription through degradation of smc5/6. Others suggest however that these incoming HBx RNA are not really required for the priming of cccDNA transcription and favor a model where HBx RNAs are the first RNA to be transcribed from established cccDNA, allowing the subsequent transcription of all the HBV viral genes^31,33^. In our study, we observed that contrary to myrcludex treated cells, in untreated cells HBx RNAs increase suggesting that they are neo-transcribed from the freshly established cccDNA. To confirm this, it will be interesting to block transcription before infection to assess the contribution of cccDNA transcription to those early HBx RNAs.

In conclusion, this novel ddOTs technique is an easy to implement and rapid assay allowing simultaneous quantification of the main HBV RNAs individually, in an accurate, reliable and highly reproducible fashion. The cost-effectiveness and speed of the procedure make it a very easy and universal way to assess the dynamics and regulation of HBV promoters, and the regulation of HBV RNA expression with unprecedented specificity. This approach may also prove useful to quantify and identify HBV circulating RNAs that could represent a powerful indicator of HBV cccDNA transcription. Finally, ddOTs approach could be adapted for the study of other organism whose genomes contain overlapping genes.

## ACKNOWLEDGEMENTS

We thank M. Benkirane for helpful discussions and critical reading of the manuscript. We thank all the members of the Molecular Virology Laboratory, Benjamin Descours and Joao Rodrigo Dias for their constructive comments. This work was supported by MSD Avenir program and the Agence Nationale de la Recherche sur le SIDA, les hépatites virales et les maladies infectieuses émergeantes (ANRS MIE). N. S. was supported by ANRS MIE and the MSD Avenir program. O. L was supported by MSD Avenir program and B. J. was supported by a fellowship from the French Ministry of Research and Technology and the ANRS MIE.

## AUTHOR CONTRIBUTIONS

1. N. S., G. P., I. M. and C. N. designed the study. N. S. performed the experiments and analyzed data, with contributions and help from B. J. and O. L.. C. N. analyzed data and wrote the manuscript with N. S.

## Competing Financial Interests

The Authors declare no competing interests

## Notes

### Competing Interest Statement

The authors have declared no competing interest.

