## supplemental figure for "ddOTs: a multiplexed quantitative ddPCR approach for resolving overlapping hepatitis B virus transcripts to decipher cccDNA-driven transcription"

Supplemental Figure 1

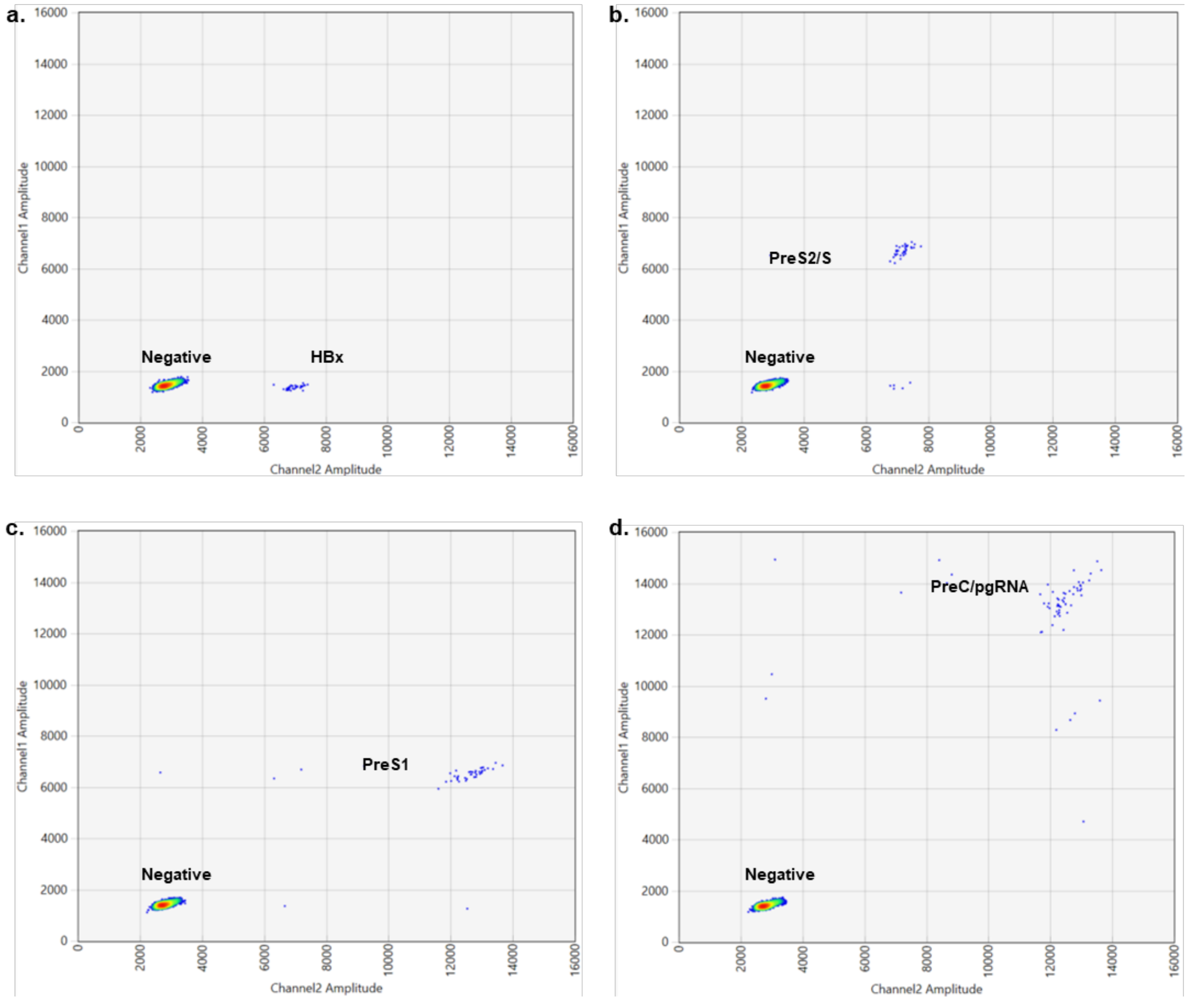


Supplemental Figure 1 : **Multiplexed ddPCR quantification of individual synthetic DNA fragments covering the 4 HBV ORFs**. Indicated PCR-amplified and purified HBV DNA fragments were quantified individually by ddOTs (a) 2D ddPCR plot of HBx DNA fragment quantification on QX Manager Software. (b) 2D ddPCR plot of synthetic PreS2/2 DNA fragment quantification on QX Manager Software. (c) 2D ddPCR plot of synthetic PreS1 DNA fragment quantification on QX Manager Software. (d) 2D ddPCR plot of synthetic preC/pregenomic DNA fragment quantification on QX Manager Software. X-axis represents HEX signal amplitude, Y-axis represents FAM signal amplitude.

Supplemental Figure 2

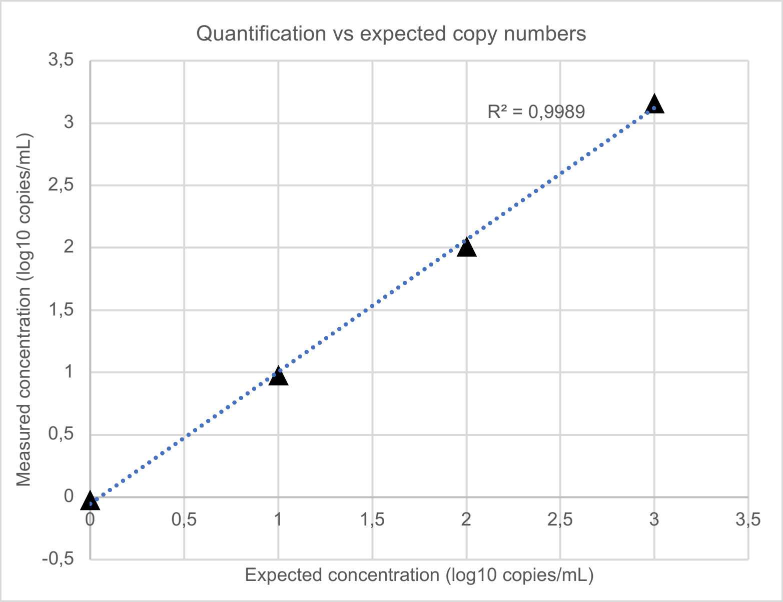


Supplemental Figure 2 : **Assessement of ddOTS linearity and sensitivity.** Correlation between known input concentrations and multiplexed ddPCR measurement of PCR-amplified and purified pregenomic DNA fragment from 1 to 1000 copy/µL.

Supplemental Figure 3

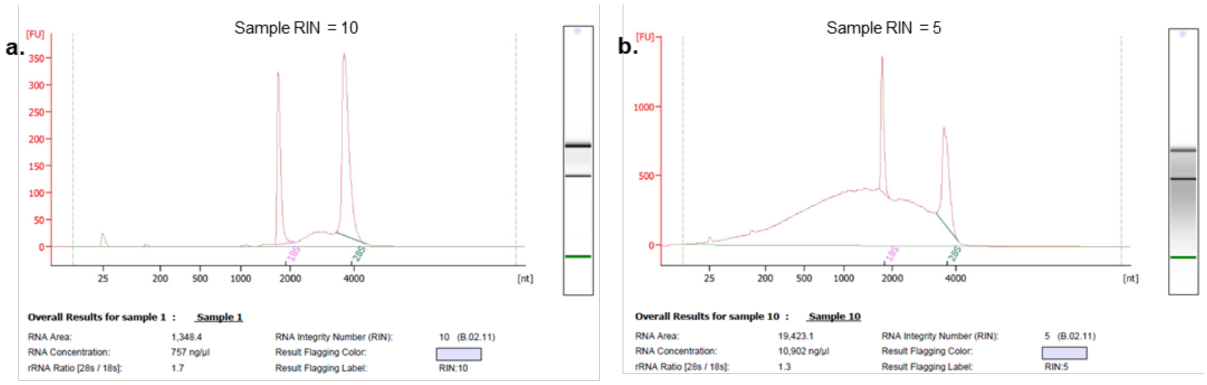


| c. | HepG2-NTCP cells infected with WT HBV RIN = 10<br>(copies/µL) | HepG2-NTCP cells infected with WT HBV RIN = 5<br>(copies/µL) |
|---|---|---|
| HBx | 26 | 108 |
| PreS2 | 122 | 112 |
| PreS1 | 43.1 | 33.1 |
| pgRNA | 71.3 | 27.6 |
| TOTAL | **262,4** | **280,7** |

Supplemental Figure 3 : **ddOTs quantification of HBV RNAs of different quality.** HepG2-NTCP cells were infected with wt HBV at a MOI of 100 vp/cells and RNA was extracted 3 days post infection. Sample was splited and one half was treated to induce degradation (temperature and vortex). (a) Bioanalyzer measurement of the RNA Integrity Number of non-treated sample which has a RIN of 10 (b) Bioanalyzer measurement of RNA Integrity Number for treated sample which has a RIN of 5. (c) Comparison of HBV RNA levels measurement of the two samples using ddOTs.
